# Limitation of External Nutrient Supply Effects by Iron-Driven Internal Phosphorus Cycling in Semi-Enclosed Coastal Waters: A Case Study of Mikawa Bay

**DOI:** 10.64898/2026.09.23.753643

**Authors:** Ippei Noda, Kiyoshi Naruse

## Abstract

The decline in fishery yields in Japan’s enclosed coastal seas has been accompanied by a reduction in phytoplankton biomass, suggesting a widespread decrease in ecosystem productivity. Oligotrophication in Mikawa Bay has caused substantial declines in phytoplankton biomass and Manila clam (Ruditapes philippinarum) production. To assess whether external nutrient enhancement could stimulate phytoplankton growth, a controlled demonstration experiment using treated effluent from two sewage treatment plants (WWTPs) was conducted (spanning fiscal years 2022 to 2024) based on analysis using a three-dimensional non-hydrostatic model. Although effluent DIP and nitrogen concentrations were increased, model results showed that WWTP-derived nutrients represent only a minor portion of the total nutrient budget. Field observations revealed limited increases in surface-water phosphorus, strong seasonal variability, and only temporary rises in chlorophyll-a and clam harvests. Monthly TP data demonstrated that seasonal DIP pulses—especially spring phosphorus bursts—dominate nutrient dynamics and overshadow external loading effects. Laboratory experiments further showed that phosphate is efficiently fixed by Fe(III) under oxidizing conditions and rapidly released during reductive dissolution of [FeOOH-HPO4]^2-^ complexes, confirming iron as the principal regulator of internal P cycling. Because current numerical models omit iron-driven internal cycling, they cannot accurately predict bioavailable P or ecosystem responses. Overall, the study indicates that localized nutrient enhancement is insufficient to counter long-term oligotrophication, and effective management must integrate nitrogen limitation, internal Fe and P cycling, oceanic exchange, and seasonal variability in Ise and Mikawa Bays.

**SYNOPSIS.:** Localized effluent nutrient enhancement failed to reverse Mikawa Bay’s oligotrophication, as internal Fe-P cycling and seasonal dynamics dominate phosphorus availability, highlighting the need for integrated, multi-factor ecosystem management.

## INTRODUCTION

In semi-enclosed coastal waters of Japan, such as Ise Bay and Mikawa Bay, marked oligotrophication has progressed in recent years due to long-term reductions in external nutrient loading as well as altered water circulation and biogeochemical processes. The resulting decline in phytoplankton biomass reduced secondary production and contributed to a significant drop in the catch of benthic organisms—particularly bivalves such as the Manila clams of Mikawa Bay (**Figure S1**).(**1–4**) Significantly, this decline initiated in the 1980s, preceding the rapid rise in sea surface temperature driven by global climate change. This timeline indicates that chronic nutrient depletion induced by water-quality management policies, rather than thermal stress, has been the primary driver of productivity loss.(**5–9**) To mitigate this oligotrophication, intentional nutrient enhancement using effluent from wastewater treatment plants (WWTPs) was explored as a potential restoration measure. However, full-scale management trials conducted from 2017 to 2024—including a two-fold increase in phosphorus concentration (Phase Ⅰ) and subsequent effluent adjustments aiming for the Redfield N:P ratio by increasing phosphorus (∼3.2×) and nitrogen (∼1.2×) loads (Phase Ⅱ)—yielded only marginal increases in phytoplankton biomass, while Manila clam landings continued to drop to historical lows (**Figure S2**).(**10–12**) These outcomes underscore the limitation of applying open-ocean stoichiometric ratios to semi-enclosed coastal ecosystems, where optimal N:P ratios vary with community structure and environmental conditions, and where nitrogen remains the primary limiting nutrient.(**13**,**14**) A fundamental obstacle to effective nutrient management is the inadequate representation of benthic-pelagic phosphorus cycling.(**15–18**) In Ise and Mikawa Bays, dissolved inorganic phosphorus (DIP) exhibits strong seasonal variability driven by sedimentary processes. Iron-mediated redox reactions regulate phosphate adsorption, release, and transformation, generating substantial springtime DIP pulses that frequently exceed external inputs.(**19**,**20**) Laboratory experiments have demonstrated the quantitative immobilization of phosphate by Fe(III) under aerobic conditions and the rapid release of phosphate accompanying the reductive dissolution of [FeOOH-HPO_4_]^2-^ complexes. Despite their importance, these internal loading mechanisms are largely omitted from current coastal ecosystem models, severely constraining their ability to predict ecological responses to external nutrient manipulations. Here, we integrate field observations, a three-dimensional nonhydrostatic hydrodynamic model, and laboratory experiments to identify the dominant controls on nutrient dynamics in Ise Bay and Mikawa Bay and evaluate the feasibility of external nutrient enhancement as a restoration strategy.

Specifically, we incorporate iron-driven internal phosphorus cycling into the framework to establish an integrated nutrient management strategy that alleviates nitrogen limitation while accounting for sediment–water phosphorus exchange. Through this approach, we aim to provide a robust scientific foundation for sustaining primary productivity and mitigating ecological risks in semi-enclosed coastal ecosystems.

## EXPERIMENTAL METHODS

### General Approach

This study was conducted as part of a multi-year field experiment in the oligotrophic Mikawa Bay to evaluate whether phytoplankton production could be increased by manipulating nutrient concentrations in WWTPs. Because the N/P ratio within phytoplankton cells tends to converge with that of the surrounding seawater, and because maximum growth in the open ocean generally occurs near the Redfield ratio (approximately 7:1 by weight), nutrient concentrations in WWTP effluents were intentionally adjusted to bring the bay-wide N/P ratio closer to this optimal range. A three-dimensional non-hydrostatic hydrodynamic model was used to assess the relative impacts of residence time, seawater exchange, and nutrient loading in Ise

Bay and Mikawa Bay.(**21–23**) Phosphorus (P) fixation in bottom sediments during winter was excluded from the model calculations because direct quantification was difficult. Simulation results indicated that increasing the P concentration in effluents from 27 WWTPs to approximately five times the usual level (2 mg/L) would significantly raise total phosphorus (TP) concentrations in both bays, potentially affecting major fishing grounds for the Manila clam.

These results demonstrate that phosphorus loading from WWTPs can have a substantial impact on nutrient dynamics in both water bodies.

### Field Experiments

Two-stage field experiments were conducted at sewage treatment plants in the Yahagi-River and Toyokawa basins to evaluate the impact of nutrient management on the coastal environment. In the first stage (FY2017 [Nov.∼Mar.], FY2018∼2019 [Oct.∼Mar.], and FY2020∼2021 [Sep.∼Mar.]), only the total phosphorus concentration in the effluent was raised to 0.76 mg/L (approximately twice the standard concentration). As no significant improvement in the fishing ground environment was observed during the first stage, the plan for the second stage involved simultaneously increasing the concentrations of total nitrogen and total phosphorus. This approach aimed to bring the N/P mass ratio within the bay closer to the Redfield ratio (N:P ratio of approximately 7:1), thereby maximizing phytoplankton production. Furthermore, the implementation period was limited to the cold season (November to April). Consequently, as shown in **Figure S3**, operations during Phase II (FY2022 [November∼March] and FY2023∼2024 [September∼March]) were conducted with target effluent concentrations set at approximately 1.3 mg/L for phosphorus (about 3.2 times the standard value) and approximately 9.6 mg/L for nitrogen (about 1.2 times the standard value). It was anticipated that during this period—despite a reduction in phosphorus bioavailability caused by the immobilization of dissolved inorganic phosphorus (DIP) by Fe(III)—it would be possible to promote spring phytoplankton blooms while minimizing the risk of red tide outbreaks.

### Environmental Monitoring and Effectiveness Evaluation

To evaluate the effectiveness of nutrient manipulation, environmental parameters including total nitrogen (TN), total phosphorus (TP), chlorophyll-a (Chl-a), dissolved oxygen (DO), water temperature, and the occurrence of red tides were monitored. Manila clam catch data were utilized as a biological indicator of trophic status alongside phytoplankton abundance. Manila clams were selected over oysters because of their high sensitivity to fluctuations in Chl-a concentration; while oysters exhibit high feeding selectivity, their extremely high filtration rates allow them to survive in low- phytoplankton environments, rendering them less representative of changes in trophic status.

Monitoring data were reviewed periodically within an adaptive management framework to adjust operational measures as necessary.

### Reagents

All chemicals were analytical-grade and used without further purification unless otherwise stated. A phosphorus standard solution (P = 1000 mg/L; Fujifilm Wako Pure Chemical Corporation) was used for all phosphate preparations. An ascorbic acid solution (72 g/L) was prepared by dissolving 7.2 g of L(+)-ascorbic acid in 100 mL of pure water. The ammonium molybdate solution was prepared by dissolving 6 g of ammonium molybdate(VI) tetrahydrate and 0.24 g of potassium antimonyl tartrate in approximately 300 mL of pure water, followed by the addition of 120 mL of sulfuric acid (2+1). The mixture was then diluted to 500 mL with pure water.

### Immobilization of Phosphate by Fe(III) under Acidic Conditions (pH < 4)

To evaluate the immobilization of phosphate by Fe(III) under weakly acidic conditions, test solutions (Samples 1∼4) were prepared as shown in **Table 1**. Standard solutions of phosphate (P = 5 mg/L) and Fe(III) (Fe = 1000 mg/L) were mixed in a stoppered test tube and diluted to the specified volume with pure water. After the mixture was allowed to stand at 20∼25°C for approximately two days, no visually observable precipitate was detected. Two milliliters of an ammonium molybdate-ascorbic acid reagent (5:1 v/v) were added to each mixture, and the reaction was allowed to proceed for 30 minutes at 20∼25°C. Subsequently, the absorbance of the supernatant was measured at 880 nm. A sample to which only 2 mL of ammonium molybdate solution had been added was used as a blank

**Table 1.**
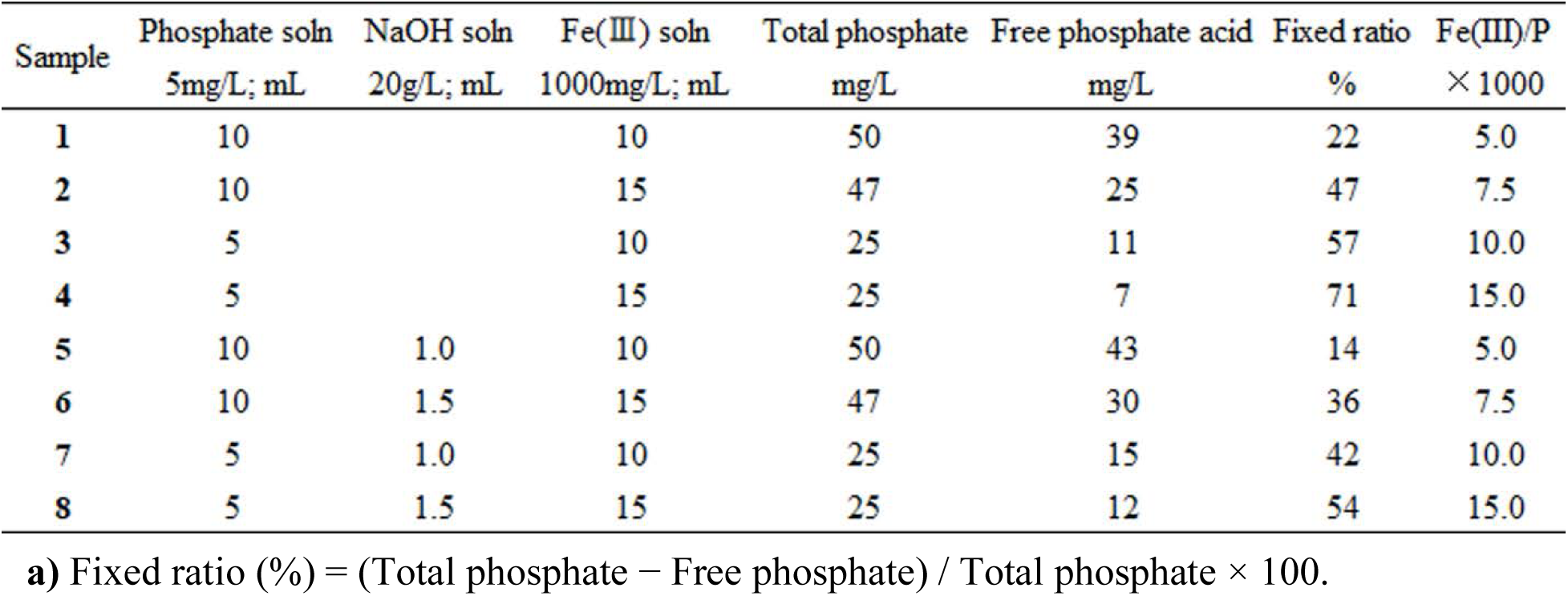
Effect of Fe(III) Addition on Dissolved Phosphate Under Acidic Conditions.

| Sample | Phosphate soln<br>5mg/L; mL | NaOH soln<br>20g/L; mL | Fe(III) soln<br>1000mg/L; mL | Total phosphate<br>mg/L | Free phosphate acid<br>mg/L | Fixed ratio<br>% | Fe(III)/P<br>× 1000 |
| --- | --- | --- | --- | --- | --- | --- | --- |
| 1 | 10 |  | 10 | 50 | 39 | 22 | 5.0 |
| 2 | 10 |  | 15 | 47 | 25 | 47 | 7.5 |
| 3 | 5 |  | 10 | 25 | 11 | 57 | 10.0 |
| 4 | 5 |  | 15 | 25 | 7 | 71 | 15.0 |
| 5 | 10 | 1.0 | 10 | 50 | 43 | 14 | 5.0 |
| 6 | 10 | 1.5 | 15 | 47 | 30 | 36 | 7.5 |
| 7 | 5 | 1.0 | 10 | 25 | 15 | 42 | 10.0 |
| 8 | 5 | 1.5 | 15 | 25 | 12 | 54 | 15.0 |

a) Fixed ratio (%) = (Total phosphate – Free phosphate) / Total phosphate × 100.

Phosphate Immobilization by [FeOOH-HPO_4_]^2-^ Complexes under Neutral Conditions (pH **≈ 8):** To simulate a closed coastal environment, phosphate immobilization under neutral conditions was evaluated using test solutions (Samples 5–8; **Table 1**). Phosphate (P = 5 mg/L) and NaOH (20 g/L) solutions were mixed, followed by the addition of Fe(III) solution (Fe = 1000 mg/L). The volume was adjusted with pure water, and the mixture was kept undisturbed at 20–25°C for 2 days. After precipitation occurred in all samples, the suspensions were centrifuged, and the supernatants were transferred to clean tubes. Each supernatant was reacted with 2 mL of the mixed reagent at 20∼25°C for 1 h before measuring absorbance at 880 nm.

Blanks were prepared using the same procedure but substituting the mixed reagent with 2 mL of ammonium molybdate solution alone.

### Phosphate Release from Iron-Phosphate Complexes under Acidic Conditions (pH < 4)

**P**hosphate release from iron-phosphate complexes under acidic conditions (pH < 4) was evaluated using Samples 9∼12 (prepared as described in **Table 2**). Phosphate (P = 5 mg/L) and Fe(III) (Fe = 1000 mg/L) solutions were sequentially mixed in a test tube, diluted to the specified volume with pure water, and left at 20∼25°C for approximately two days. Next, an alkaline NaBH₄ solution (10 g/L, 0.5 mL) was added dropwise under stirring, and the mixture was reacted at 20–25°C for 1 day. After adding 2 mL of the mixed reagent (5:1 v/v), the solution was reacted for another 30 minutes before measuring absorbance at 880 nm. Blanks were prepared identically using ammonium molybdate solution (2 mL) alone.

**Table 2.**
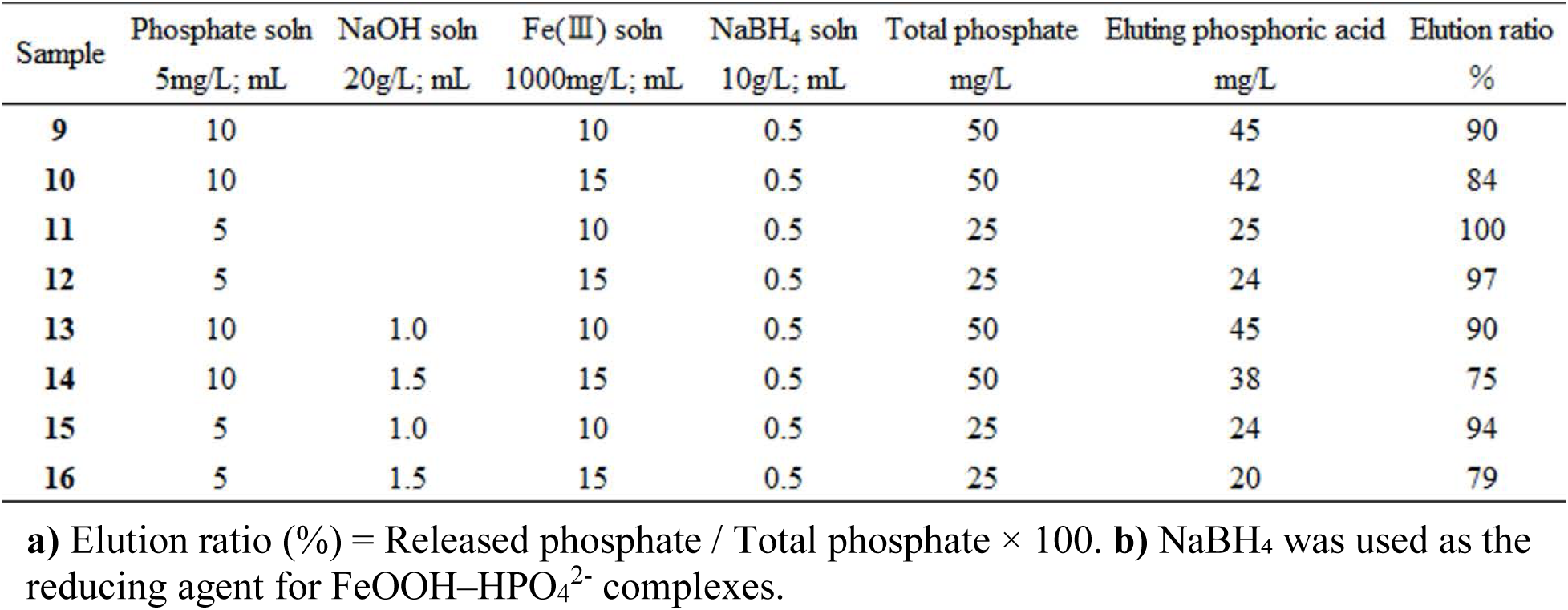
Dissolution of Fe(III) and Release of Phosphate Accompanying the Reduction of [FeOOH-HPO_4_]^2-^ Complexes.

| Sample | Phosphate soln<br>5mg/L; mL | NaOH soln<br>20g/L; mL | Fe(III) soln<br>1000mg/L; mL | NaBH<sub>4</sub> soln<br>10g/L; mL | Total phosphate<br>mg/L | Eluting phosphoric acid<br>mg/L | Elution ratio<br>% |
| --- | --- | --- | --- | --- | --- | --- | --- |
| 9 | 10 |  | 10 | 0.5 | 50 | 45 | 90 |
| 10 | 10 |  | 15 | 0.5 | 50 | 42 | 84 |
| 11 | 5 |  | 10 | 0.5 | 25 | 25 | 100 |
| 12 | 5 |  | 15 | 0.5 | 25 | 24 | 97 |
| 13 | 10 | 1.0 | 10 | 0.5 | 50 | 45 | 90 |
| 14 | 10 | 1.5 | 15 | 0.5 | 50 | 38 | 75 |
| 15 | 5 | 1.0 | 10 | 0.5 | 25 | 24 | 94 |
| 16 | 5 | 1.5 | 15 | 0.5 | 25 | 20 | 79 |

**a)** Elution ratio (%) = Released phosphate / Total phosphate × 100. **b)** NaBH<sub>4</sub> was used as the reducing agent for FeOOH-HPO<sub>4</sub><sup>2-</sup> complexes.

### Phosphate Release from [FeOOH-HPO4]^2-^ Complexes under Neutral Conditions (pH ≈ 8)

Phosphate release under neutral conditions was evaluated using Samples 13∼16 (prepared as described in **Table 2**). A phosphate solution (P = 5 mg/L) and an NaOH solution (20 g/L) were mixed, followed by the addition of an Fe(III) solution (Fe = 1000 mg/L). After adjusting the volume with pure water and homogenizing, the mixture was allowed to stand at 20∼25°C for approximately 2 days. Next, an alkaline NaBH₄ solution (10 g/L, 0.5 mL) was added, and the solution was reacted at 20∼25°C for 1 day. Subsequently, 2 mL of the mixed reagent (5:1 v/v) was added, and after reacting for 1 hour, the absorbance at 880 nm was measured. Blanks were prepared identically using only ammonium molybdate solution (2 mL).

## RESULTS AND DISCUSSION

Recent oligotrophication in Mikawa Bay has caused a marked decline in phytoplankton biomass, leading to a significant drop in the catch of benthic organisms such as Manila clams. This study evaluated whether phytoplankton growth can be stimulated by manipulating nutrient concentrations in effluent from WWTPs. A three-dimensional non-hydrostatic model was first applied to quantify residence time, oceanic exchange, and the relative contributions of external nutrient loads in Ise and Mikawa Bays. Although the model does not explicitly represent seasonal P cycling, it indicates that WWTP-derived nutrient inputs constitute only a minor fraction of the total nutrient budget.

### Dominant Controls on Nutrient Dynamics and the Limitations of External Loading Manipulation

Guided by the model results, a controlled field experiment was conducted from fiscal years 2022 to 2024 in which nutrient concentrations in effluent from the Yahagi-River and Toyokawa WWTPs were intentionally modified. DIP concentrations were increased to approximately 3.2 times the normal level, and dissolved nitrogen concentrations were increased by roughly 1.2 times. Long-term monitoring data for surface-water nutrients and Chl-a at stations near the WWTPs were used to assess the ecological response. As shown in **Figure 1a**, a significant increase in TP was observed only at monitoring point K-7, located adjacent to the Yahagi-River WWTP. This is consistent with the fact that the discharge volume of the Yahagi-River WWTP (277,560 m³/day) is larger than that of the Toyokawa WWTP (91,879 m³/day). In contrast, total TN at K-7 showed little change, reflecting the smaller increase in N relative to P (**Figure 1b**). Chl-a at K-7 increased temporarily in FY2022 but subsequently declined (**Figure 1c**), and Manila clam harvests exhibited a similar pattern; an initial increase followed by a sustained decline to record-low levels. These results reveal a pronounced nonlinearity between external nutrient enhancement and ecosystem response. To resolve this discrepancy, monthly TP data were analyzed to examine seasonal DIP dynamics. **Figure 1d** showed that, despite the increase in DIP, surface water TP did not consistently increase and, during certain periods, fell below the levels observed during normal operation. The Chl-a increase in FY2022 was attributable not to the DIP manipulation but to a spring P-bloom that occurred prior to the experiment. These findings indicate that surface-water P concentrations are governed primarily by internal cycling processes including redox-driven transformations associated with the oxidation of organic and ammonium nitrogen in bottom waters, sediment resuspension and release, and seasonal stratification and mixing—rather than by external loads. The timing of the spring phytoplankton burst strongly determines food availability for Manila clam and directly influences harvest outcomes. Overall, seasonal DIP variability overwhelms the effects of external DIP enhancement. DIP concentrations do not necessarily increase during manipulation periods and may even fall below baseline levels. The observed delay of the spring bloom and its shift toward a summer bloom can be partially attributed to reduced microbial activity caused by the depletion of organic and ammonium nitrogen in bottom waters, which suppresses internal nutrient recycling. Although internal P cycling was not explicitly represented in the model, the model results and observations together indicate that oceanic exchange and river discharge are the dominant controls on nutrient dynamics in Mikawa Bay. The magnitude of nutrient enhancement implemented in the field experiment was insufficient to offset the long-term decline in nutrient concentrations. Seasonal P pulses, such as the spring DIP burst, likely counteracted the effects of external loading. Regression analysis of the relationships between P and Chl-a and between N and Chl-a confirmed that Ise Bay and Mikawa Bay are systems rich in P but limited by N, and that N is the primary factor constraining phytoplankton growth. Multivariate analyses further demonstrated that both N/P ratios and absolute nutrient concentrations influence Chl-a.(**15**) Notably, **Figure S3** shows that increasing P and N in accordance with the Redfield ratio in oligotrophic coastal waters can lead to a decrease in phytoplankton biomass, suggesting that the Redfield ratio is not necessarily optimal for that environment. Seasonal differences in ecosystem response were also evident. **Figure 2** shows that while a rapid increase in P triggered a surge in Chl-a during spring, no such response was observed in late summer or autumn. Despite efforts to increase winter TP concentrations from FY2017 to FY2025, spring DIP bursts did not occur, suggesting strengthened P retention in sediments. A plausible mechanism is reduced activity of iron-reducing bacteria due to early winter N depletion, which suppresses DIP release from sediments and may delay P increases until summer.(**24–26**)

**Figure 1.**
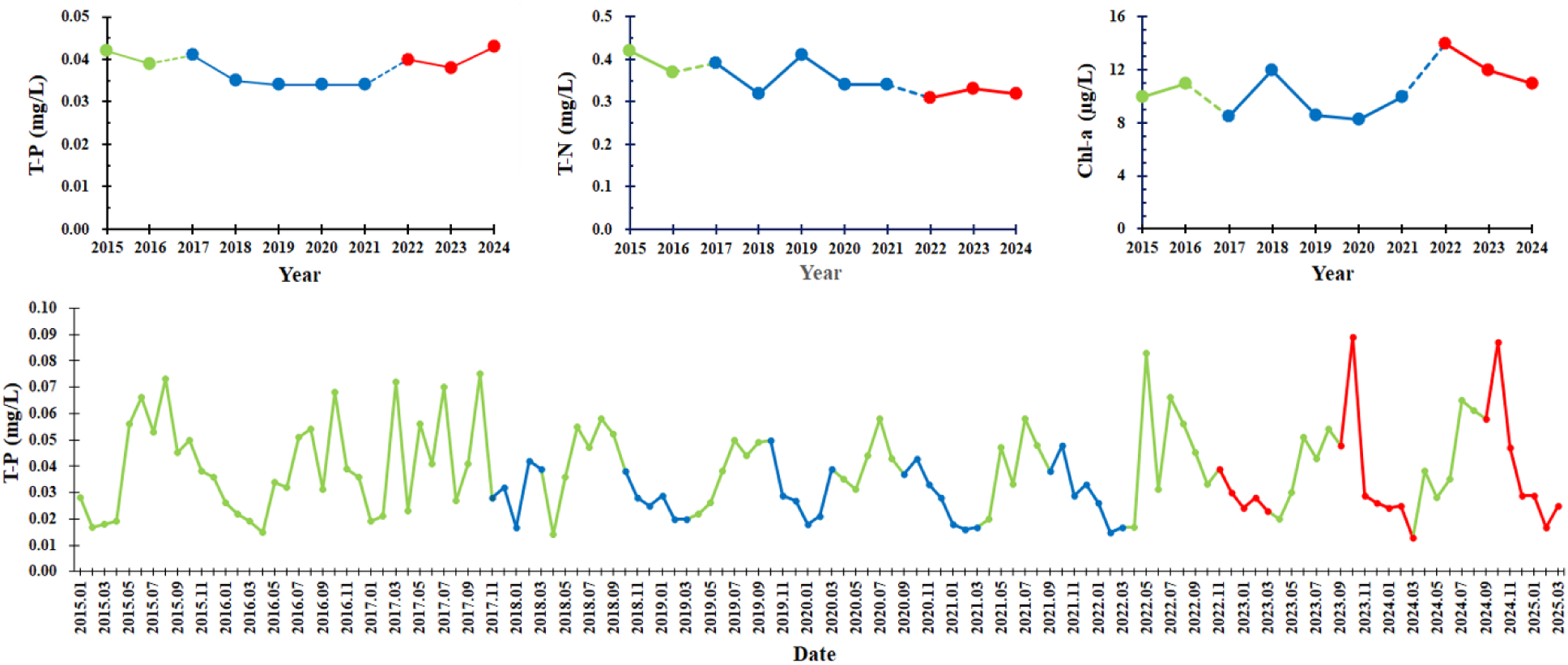
Seasonal variations in TP (surface DIP), TN, and Chl-a at observation point K-7 under conditions of increased external P loading. Despite the increase in P loading from WWTP effluent, surface P concentrations did not rise during the experimental period. Instead, a notable feature was a rapid increase (burst) in DIP during the spring. These results suggest that seasonal variations in P outweigh the influence of external loading. Variations in surface water quality at K-7. (**a**–**c**) Inter- annual variations in surface TP (**a**), TN (**b**), and Chl-a (**c**); (**d**) long-term monthly variation in surface TP. Symbols indicate levels relative to normal operation: —Phase I (TP ∼2×) and —Phase II (TP ∼3.2×, TN ∼1.2×).

**Figure 2.**
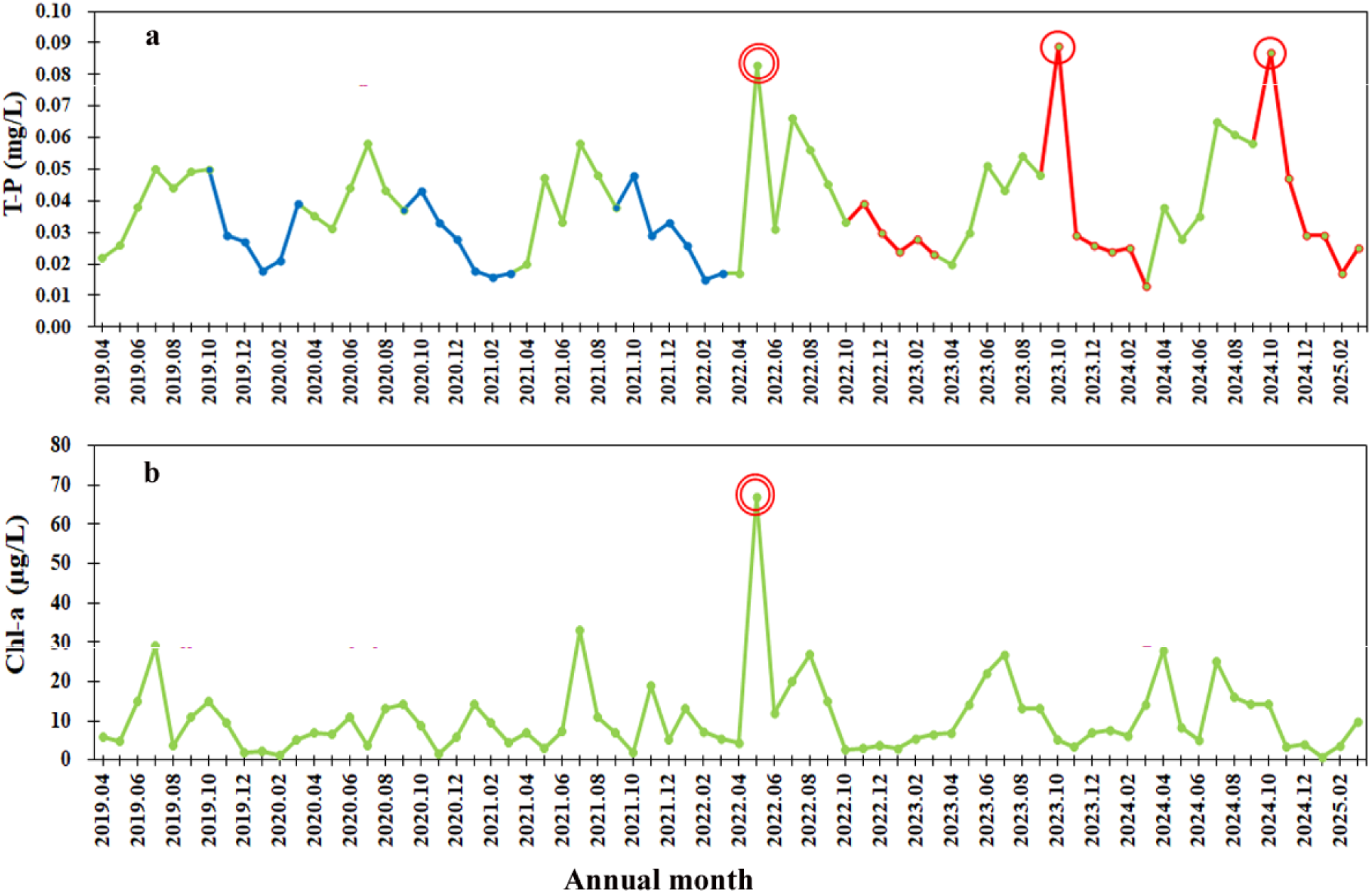
Differences in ecosystem responses to seasonal P surges. This figure shows that while a spring P surge leads to an increase in Chl-a, a similar response is not observed from late summer to autumn; furthermore, interventions that raised winter TP concentrations did not result in a spring DIP surge (burst). **a.** Interventions to increase winter TP concentrations. — indicates a 3.2-fold increase (Phase II), — indicates a 2-fold increase (Phase I). Spring P surge: ○; late summer to autumn P surge: ○. **b.** Spring surge in Chl-a concentration: ○.

When both P and N loads were increased—rather than P alone—Chl-a increases were observed, consistent with the temporary rise in Manila clam harvests in FY2022. However, harvests declined thereafter, reaching historically low levels. These results collectively underscore the importance of understanding and managing nitrogen dynamics in Ise and Mikawa Bays. Because physical processes such as oceanic exchange and seasonal mixing exert strong control over nutrient concentrations, localized load manipulation alone has limited capacity to induce desired ecosystem responses. Future nutrient-management strategies must integrate external loads, internal cycling, oceanic exchange, and seasonal variability.

### Redox-Driven Seasonal Cycling of Phosphorus in Mikawa Bay and Ise Bay

The seasonal dynamics of iron (Fe) and phosphorus (P) in Mikawa Bay and Ise Bay are shaped by biogeochemical processes characteristic of semi-enclosed coastal systems, including riverine inputs, surface-water chemical reactions, sedimentary retention, and benthic internal loading. These processes exhibit a consistent annual pattern in which the redox cycling of iron exerts dominant control over phosphorus mobility. P in surface waters is removed from the dissolved phase through reactions with Fe^3+^ and Al^3+^ supplied by rivers (**Figure 3**) and with Ca^2+^, which is abundant in seawater. These cations form sparingly soluble phosphate minerals or strong adsorption complexes that immobilize P. Under acidic conditions, Fe^3+^ and Al^3+^ interact mainly with protonated phosphate species (H_2_PO_4_ ^-^ and, to a lesser extent, HPO_4_ ^2-^). Owing to the lower solubility of Fe-phosphate minerals relative to Al–phosphate minerals, Fe^3+^ preferentially immobilizes dissolved phosphorus when both cations coexist.(**27**) Under alkaline conditions, Ca^2+^ readily forms low-solubility calcium phosphate phases. In seawater at pH ∼8, Al occurs predominantly as the tetrahydroxo complex Al(OH) ^-^, which exhibits negligible affinity for HPO_4_ ^2-^. In contrast, HPO_4_ ^2-^ effectively immobilizes phosphate by binding strongly to Fe(III) oxyhydroxides (FeOOH)—particularly goethite and ferrihydrite—to form stable inner-sphere surface complexes.(**28**) Consequently, Fe(III) phases serve as the principal P-fixing agents in the surface waters of both bays. In bottom waters, internal loading and reductive release of P occur during summer. Stratification and elevated temperatures reduce dissolved oxygen (DO), driving sediments toward strongly reducing conditions. Under such conditions, the stability of P-bearing phases diverges markedly. Calcium phosphate [Ca_3_(PO_4_)_2_] is largely insensitive to redox changes and releases minimal DIP. In contrast, [FeOOH-HPO_4_]^2-^ undergoes reductive dissolution as Fe(III) is converted to Fe(II), liberating HPO_4_ ^2-^ into bottom waters. Al-bound phosphate is more stable than Fe-bound phases and exhibits limited reductive dissolution. Microbial Fe reduction, fueled by oxidation of organic matter and sulfides, accelerates Fe^3+^→Fe^2+^ conversion and consumes DO.(**28**,**29**) As a result, hypoxia and rapid DIP accumulation proceed simultaneously. The formation of summer hypoxic layers and episodic “blue tides” in the region is attributed to this chain reaction of Fe reduction and P release. Seasonal redox cycling produces a characteristic annual pattern in P dynamics. In winter, full-water-column oxygenation promotes oxidation of Fe^2+^ to Fe^3+^ and reformation of FeOOH, enabling renewed adsorption of HPO_4_ ^2-^ and sedimentary P fixation. In summer, declining DO in bottom waters reduces [FeOOH-HPO_4_]^2-^ and releases P. The resulting Fe^2+^ reacts with HS^-^ produced by sulfate-reducing bacteria(**29,30**) to form FeS, suppressing re-fixation of P by Fe.(**31**) In autumn, cooling and vertical mixing restore oxygen to bottom waters, oxidizing Fe^2+^ to Fe^3+^ and enabling re-immobilization of P as [FeOOH-HPO_4_]^2-^. DIP released from sediments during summer is transported upward during winter mixing and serves as a major nutrient source for the spring phytoplankton bloom. Collectively, these observations indicate that P fixation and its seasonal variability in both bays are governed primarily by the redox-driven cycling of Fe.

**Figure 3.**
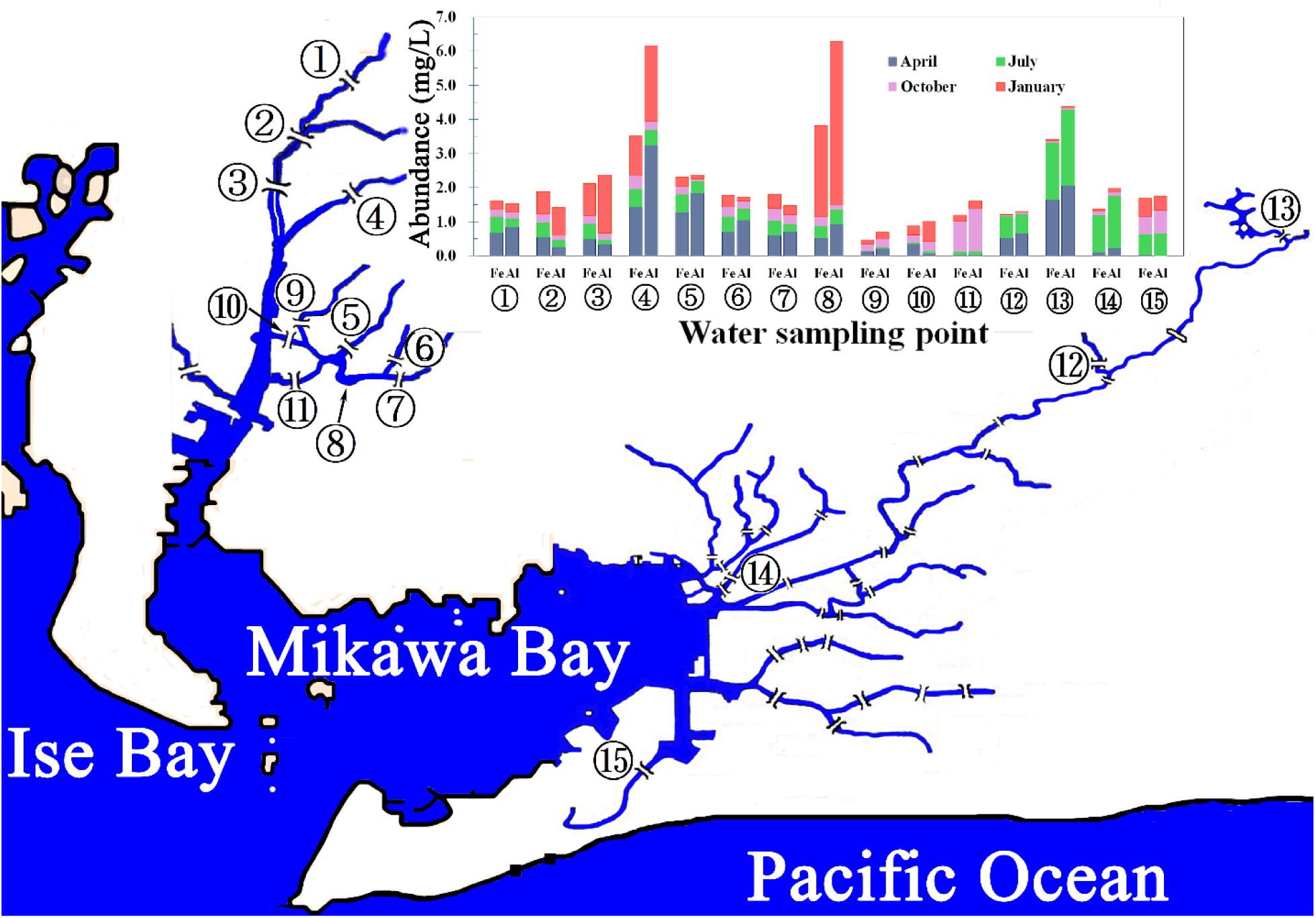
The iron and aluminum contents in river water samples collected quarterly at observation points (1) through (15) during the 2021 fiscal year were measured using ICP-MS. (1): Shinkai- bashi, (2): Sakai-gawa, (3): Ichihara-bashi, (4): Aizuma-gawa, (5): Tansui-bashi, (6): Sakashita-bashi, (7): Sakashita-kobashi, (8): Aburagafuchi, (9): Hieda-bashi, (10): Takahama-bashi, (11): Suimon-bashi, (12): Nagashino-bashi, (13): Horai-bashi, (14): Yanagi-bashi, (15): Funakura-bashi.

### Iron-Mediated Phosphorus Cycling and Limitations of Current Numerical Models; Iron Redox Reactions and P Desorption at the Sediment–Water Interface

his study tested the hypothesis that iron acts as the principal regulatory factor controlling internal P cycling— specifically the fixation and release processes occurring at the sediment–water interface—in Ise Bay and Mikawa Bay. Because iron readily undergoes rapid redox transitions between Fe(II) and Fe(III) at relatively low redox potentials, it responds sensitively to environmental fluctuations. This chemical property makes iron a key determinant of phosphate adsorption–desorption dynamics in coastal sediments. In the water column, Eh values in surface and mid-depth layers range from 450 to 150 mV, indicating oxidizing conditions. In contrast, summer sediments exhibit Eh values declining to 150–0 mV, and bottom waters reach strongly reducing conditions (−300 to −100 mV).(**32–35**) Under such conditions, terrestrially derived Fe(III) is microbially reduced to Fe(II), increasing the likelihood that phosphate previously bound to Fe(III) phases will be released into porewaters and subsequently into overlying bottom waters.

### Quantitative Evaluation of Phosphate Immobilization Under Oxidizing Conditions

To elucidate phosphate fixation mechanisms under oxidizing conditions, FeCl₃·6H₂O and KH₂PO₄ were reacted at pH < 4, and dissolved phosphate concentrations were monitored over time. No solid FePO₄ precipitates were visually or analytically detected, indicating that excess Fe(III) remained primarily as soluble Fe^3+^. Nevertheless, dissolved phosphate concentrations decreased proportionally with Fe(III) additions (**Table 1**, samples 1–4), demonstrating quantitative fixation without phase transformation. This suggests that phosphate was immobilized through formation of inner-sphere complexes (e.g., Fe–H_2_PO_4_ ^+^) or ion pairs in solution, consistent with the high solubility of FePO_4_ under acidic conditions.(**36**) At seawater pH (∼8), Fe(III) exists mainly as FeOOH and phosphate as HPO_4_^2-^, forming [FeOOH-HPO_4_]^2-^ composite precipitates.(**37–40**)

Excess Fe(Ⅲ) precipitates as FeOOH. As shown in **Table 1** and **Figure 4**, dissolved phosphate decreased linearly with increasing Fe(Ⅲ) additions, revealing a positive linear relationship between the Fe(Ⅲ)/P ratio and phosphate-fixation efficiency. Free Fe³⁺ exhibited higher fixation capacity than FeOOH, but substantial FeOOH quantities were required to effectively remove low concentrations of phosphate typical of seawater.

**Figure 4.**
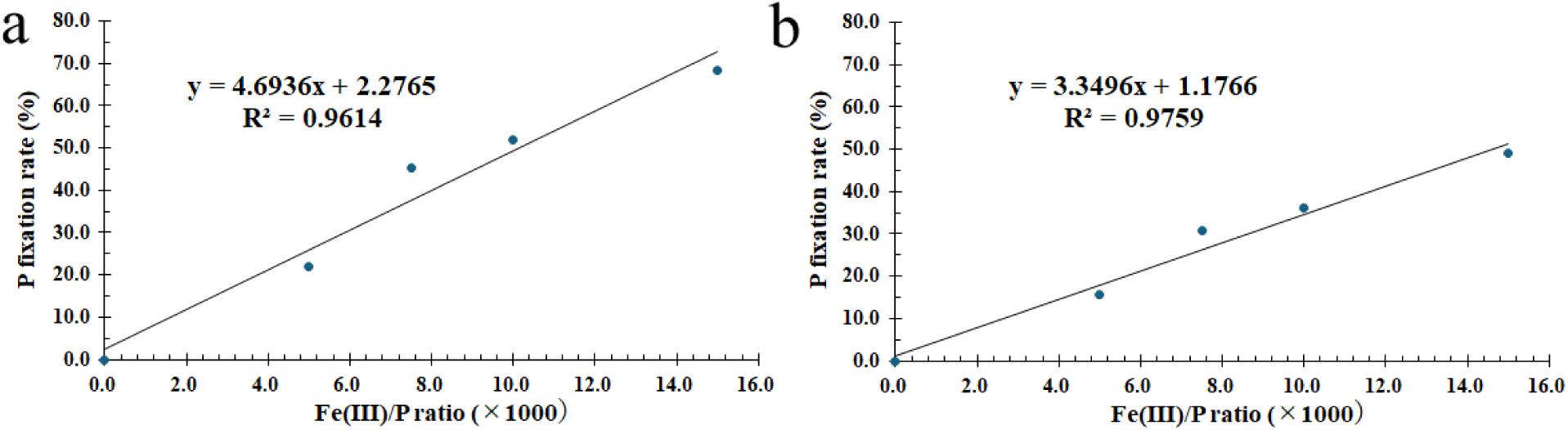
Linear relationship between the Fe(Ⅲ)/P ratio and the amount of immobilized phosphate: This demonstrates that dissolved phosphate decreases linearly as the amount of added Fe(Ⅲ) increases, serving as a quantitative indicator of immobilization efficiency. **a**: Acidic conditions. **b**: Neutral conditions.

### Phosphate Release Under Reducing Conditions and Implications for Numerical Modeling

To simulate reductive release, synthetic [FeOOH-HPO_4_]^2-^ complexes were chemically reduced using NaBH₄. As shown in **Table 2**, Fe(Ⅲ) was rapidly reduced to Fe(Ⅱ), accompanied by dissolution and release of phosphate into solution. This experiment mimics natural microbial reduction of FeOOH and demonstrates that Fe(Ⅲ) in both FeOOH and [FeOOH-HPO4]^2-^ dissolves non-selectively under reducing conditions. These findings provide experimental evidence that iron redox cycling is the primary driver of P immobilization, release, and seasonal variability in Ise Bay and Mikawa Bay. They also highlight the structural limitations of the "three-dimensional non-hydrostatic model" used in recent field experiments, which failed to predict substantial seasonal increases in phytoplankton biomass. Specifically, the model did not incorporate iron-driven internal P cycling, treating DIP dynamics primarily as functions of external loading and hydrodynamic transport. Specifically, the model did not incorporate iron-driven internal P cycling and treated DIP dynamics mainly as functions of external loading and hydrodynamic transport. As demonstrated here, the quantities of Fe³⁺, ambient phosphate concentrations, and Fe/P ratios are critical parameters determining phosphate-fixation efficiency. Because these quantitative relationships had not been formulated previously, earlier models lacked essential parameters for predicting the bioavailable fraction of added phosphorus. By experimentally resolving the quantitative relationship between Fe³⁺ and phosphate fixation under environmentally relevant conditions, this study fills a methodological gap in coastal biogeochemical modeling. The results provide essential knowledge for advancing future modeling of Ise Bay and Mikawa Bay and for developing effective nutrient-management strategies.

### The Necessity of "Integrated Nutrient Management" for the Recovery of Coastal Marine Resources

To restore coastal marine resources, management strategies must shift from targeting external nutrient input loads to implementing "Integrated Nutrient Management," which links sediment geochemistry and microbial processes (specifically the iron-phosphorus cycles) with biological responses such as N limitation and Manila clam production. In N-limited estuaries like Ise and Mikawa Bays, early-winter N deficiencies (both organic and ammonium forms) suppress the activity of iron-reducing bacteria, which, combined with internal cycling dynamics, inhibits the release of DIP from sediments. Consequently, simply supplying P fails to trigger the rapid spring surge (pulse) in DIP concentration; instead, it promotes sediment phosphorus accumulation and delays phytoplankton blooms until summer. This indicates that despite their phosphorus abundance, primary production in these bays is strictly governed by N availability. The temporary increase in Manila clam landings during fiscal year 2022, following a simultaneous rise in N and P loads, supports the idea that nitrogen input improves food availability. However, minor load adjustments or uniform nutrient additions based solely on the Redfield ratio cannot overcome physical oceanographic transport—such as vertical mixing and exchange with open-ocean waters—to achieve long-term resource recovery. Future management requires large-scale numerical models integrating hydrodynamics and ecosystem dynamics. Such frameworks must incorporate optimized, site-specific N/P ratios and targeted early-winter nitrogen inputs to sustain bottom-water microbial processes and ensure ecosystem restoration.

## CONCLUTION

This study demonstrates that increasing nutrient concentrations based on the Redfield ratio of sewage treatment plant effluent cannot reverse the long-term oligotrophication of Mikawa Bay. External DIP and nitrogen additions produced only minor and temporary ecological responses because nutrient dynamics are dominated by oceanic exchange, seasonal stratification–mixing, and redox-driven internal phosphorus cycling. Laboratory experiments confirmed that iron governs phosphate fixation and release, a key mechanism absent from current numerical models. As a result, external loading alone cannot reliably enhance phytoplankton biomass or support sustained Manila clam recovery. Effective nutrient-management strategies must integrate nitrogen limitation, Fe and P cycling, hydrodynamic processes, and seasonal variability to accurately predict bioavailable nutrients and restore ecosystem productivity in Ise and Mikawa Bays.

## ASSOCIATED CONTENTS

### Supporting Information

Long-term trends in Manila clam catches in Japanese coastal waters. (**Figure S1**); Long-term trends in Manila clam catches in Aichi Prefecture. (**Figure S2**); Operational conditions for sewage treatment plant effluent in the Phase II demonstration test. (**Figure S3**); Analysis of Phytoplankton Response Integrating Nutrient Concentrations and N/P Ratios. **(Figure S4)**; Differences in Phosphate Immobilization Mechanisms Depending on Reaction Conditions. (**Figures S3, S4, S5**)

## AUTHOR INFORMATION

Corresponding Author

## Author

Kiyoshi Naruse

## Author Contributions

I.N. conceived, designed, and supervised the study and conducted the sampling. K.N. raised questions and issues regarding the manuscript from an ecological perspective, and I.N. revised the manuscript accordingly. All authors contributed to writing the manuscript and approved the final version.

## Notes

The author declares no competing financial interest.

## ACKNOWLEDGMENT

This research result is part of the results of a survey conducted by Aichi Prefecture in Mikawa Bay and Ise Bay, and we would like to thank the relevant staff for their helpful advice. We thank Takanori Matui, Tarou Okuno, Norimasa Naruse, Seigo Horie, Tadashi Endou, Masanari Miwa, Yoshihiro Yokoi (Bureau of the Environment, Aichi Pref.) for assistance in the field. We thank Professor Toshihiko Yokoyama (Institute for Molecular Science) for helpful discussions.

## Notes

### Competing Interest Statement

The authors have declared no competing interest.

