## Supplemental Figure S1-S7 for "Limitation of External Nutrient Supply Effects by Iron-Driven Internal Phosphorus Cycling in Semi-Enclosed Coastal Waters: A Case Study of Mikawa Bay"

### Support Information

### Contents

1. Long-term trends in Manila clam catches in Japanese coastal waters. (**Figure S1**)

・・・・・pages 2

2. Long-term trends in Manila clam catches in Aichi Prefecture. (**Figure S2**)

・・・・・pages 2~3

3. Operational conditions for sewage treatment plant effluent in the Phase II demonstration test. (**Figure S3**)

・・・・・pages 3~4

4. **Analysis of Phytoplankton Response Integrating Nutrient Concentrations and N/P Ratios.** (**Figure S4**)

・・・・・pages 4~5

5. **Differences in Phosphate Immobilization Mechanisms Depending on Reaction Conditions.** (**Figure S5~S7**)

・・・・・pages 5~6

#### 1. Long-term trends in Manila clam catches in Japanese coastal waters.

During Japan's period of rapid economic growth, eutrophication in enclosed coastal waters—caused by land-based pollution—led to frequent red tides and blue tides (hypoxic water upwelling), resulting in mass mortality of marine life and severe damage to the fishing industry. In response, coastal municipalities implemented rigorous measures against red tides. Subsequently, a marked decline in fishery catches was observed in these enclosed waters. This decline coincided with a reduction in phytoplankton biomass, suggesting a drop in the productivity of the entire ecosystem. This trend is believed to stem primarily from the long-term reduction of nitrogen and phosphorus loads—implemented as a countermeasure against red tides—whereby a decrease in nutrient supply led to reduced primary production. Furthermore, as shown in **Figure S1**, alongside this decline in productivity, populations of the Manila clam—a species highly sensitive to phytoplankton levels—began to show a downward trend nationwide around 1984. This decline in productivity became apparent in the 1990s, predating the rise in sea surface temperatures associated with global warming. Therefore, the primary cause is considered to be not temperature change, but a state of "chronic nutrient deficiency (oligotrophication)" resulting from excessive water quality improvement measures.

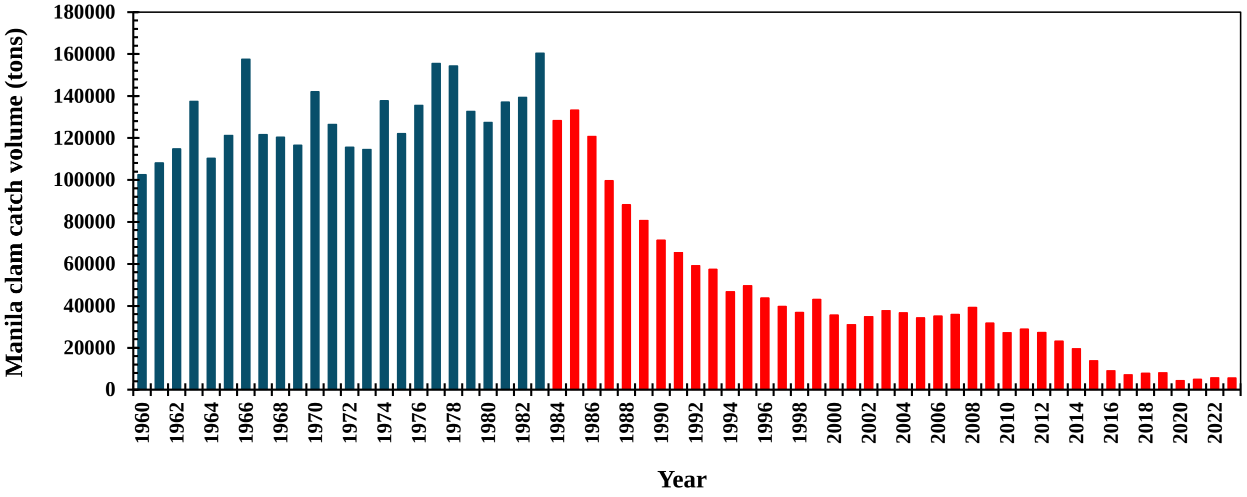


**Figure S1.** Trends in the annual catch of Manila clams in Japan. Although some improvement was observed following the release of juvenile clams, the catch has declined sharply since fiscal year 1984, repeatedly hitting new record lows Catch levels prior to fiscal year 1984 and the downward trend in catches from fiscal year 1984 onwards.

#### 2. Long-term trends in Manila clam catches in Aichi Prefecture.

Aichi Prefecture (particularly Mikawa Bay) had held the top spot in Japan for Manila clam catches since 2008 but fell to second place in 2024. To address these ecosystem changes, initiatives are underway to restore primary production by intentionally supplying nutrients (nutrient supplementation), utilizing treated wastewater from sewage treatment plants. These measures aim to increase phytoplankton in nutrient-deficient coastal areas by replenishing nutrients that would otherwise be supplied from rivers or human activities.

Furthermore, as shown in **Figure S2**, the decline in Manila clam catches in Aichi Prefecture began in the 2010s. This occurred significantly later than the declines observed in other coastal

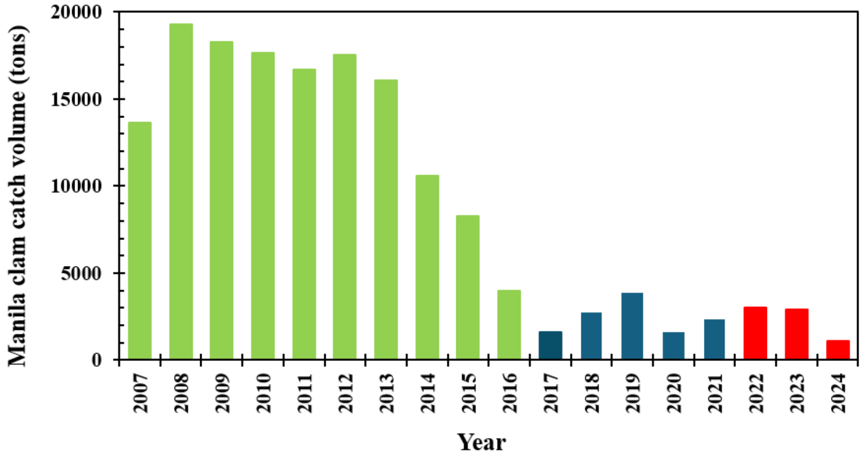


**Figure S2.** Trends in annual Manila clam catches in Aichi Prefecture. Although some improvement was observed following the release of juvenile clams, catches have plummeted since the 2013 fiscal year, repeatedly hitting new record lows. The delay in the decline of catches compared to other regions across Japan is attributed to the fact that regulations regarding pollution loads were implemented later than in areas such as the Seto Inland Sea and Tokyo Bay. ▬ : Phase II; ▬ : Phase I.

regions of Japan, such as the Seto Inland Sea and Tokyo Bay, where the decrease had already begun in the 1980s. Located in central Japan—specifically along the Pacific coast—Aichi Prefecture serves as a key indicator region for understanding the impacts of climate change and is crucial for assessing how climate change affects the country as a whole. This delayed onset is believed to stem from the fact that Aichi Prefecture implemented water quality regulations later than other regions. These trends suggest that the primary cause of both the decline in phytoplankton and the drop in Manila clam catches is the oligotrophication (nutrient depletion) of this semi-enclosed sea area.

#### 3. Operational conditions for sewage treatment plant effluent in the Phase II demonstration test.

Based on numerical model simulations, the regulatory standards for TP and total nitrogen (TN) were relaxed in fiscal year 2022, increasing from 1.0 mg/L to 2.0 mg/L and from 10 mg/L to 20 mg/L, respectively. Accordingly, as shown in **Figure S3**, operations during Phase II (November–

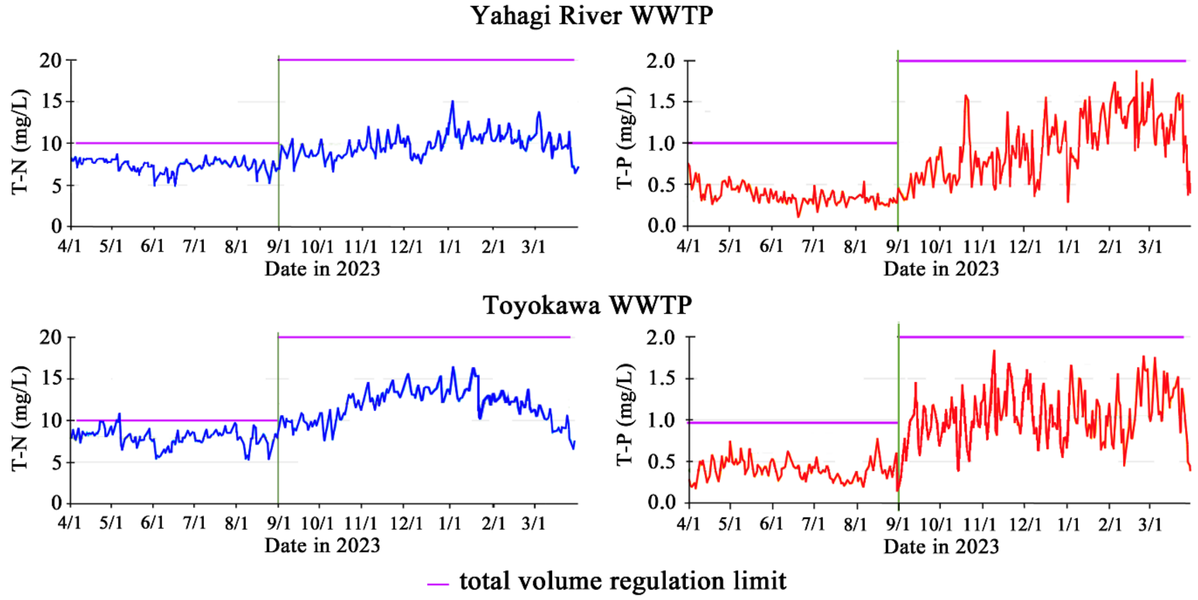


**Figure S3.** Changes in dissolved inorganic phosphorus (DIP) and dissolved nitrogen concentrations in effluent from the Yahagi-River WWTP and Toyogawa WWTP during the increased-load operation period and the normal operation period in fiscal year 2023.

March of fiscal year 2022, and September–March of fiscal years 2023–2024) were conducted with target effluent concentrations set at approximately 1.3 mg/L for phosphorus (about 3.2 times the standard value) and approximately 9.6 mg/L for nitrogen (about 1.2 times the standard value).

#### 4. Analysis of Phytoplankton Response Integrating Nutrient Concentrations and N/P Ratios.

Recent studies have revealed that the nutrient requirements of phytoplankton vary depending on the trophic status of the water body and that the N/P ratio plays a crucial role in mitigating the effects of oligotrophication. However, conventional analyses linking the N/P ratio to chlorophyll-a (Chl-a) concentrations often overlook the influence of absolute nutrient quantities, potentially leading to misinterpretations. In reality, phytoplankton growth is likely governed more by the total amount of nutrients than by their relative ratios. Therefore, this study employed multivariate analysis to evaluate the combined effects of the N/P ratio and absolute nutrient concentrations on Chl-a levels. An assessment of measures implemented during "Phase II" using this approach revealed that

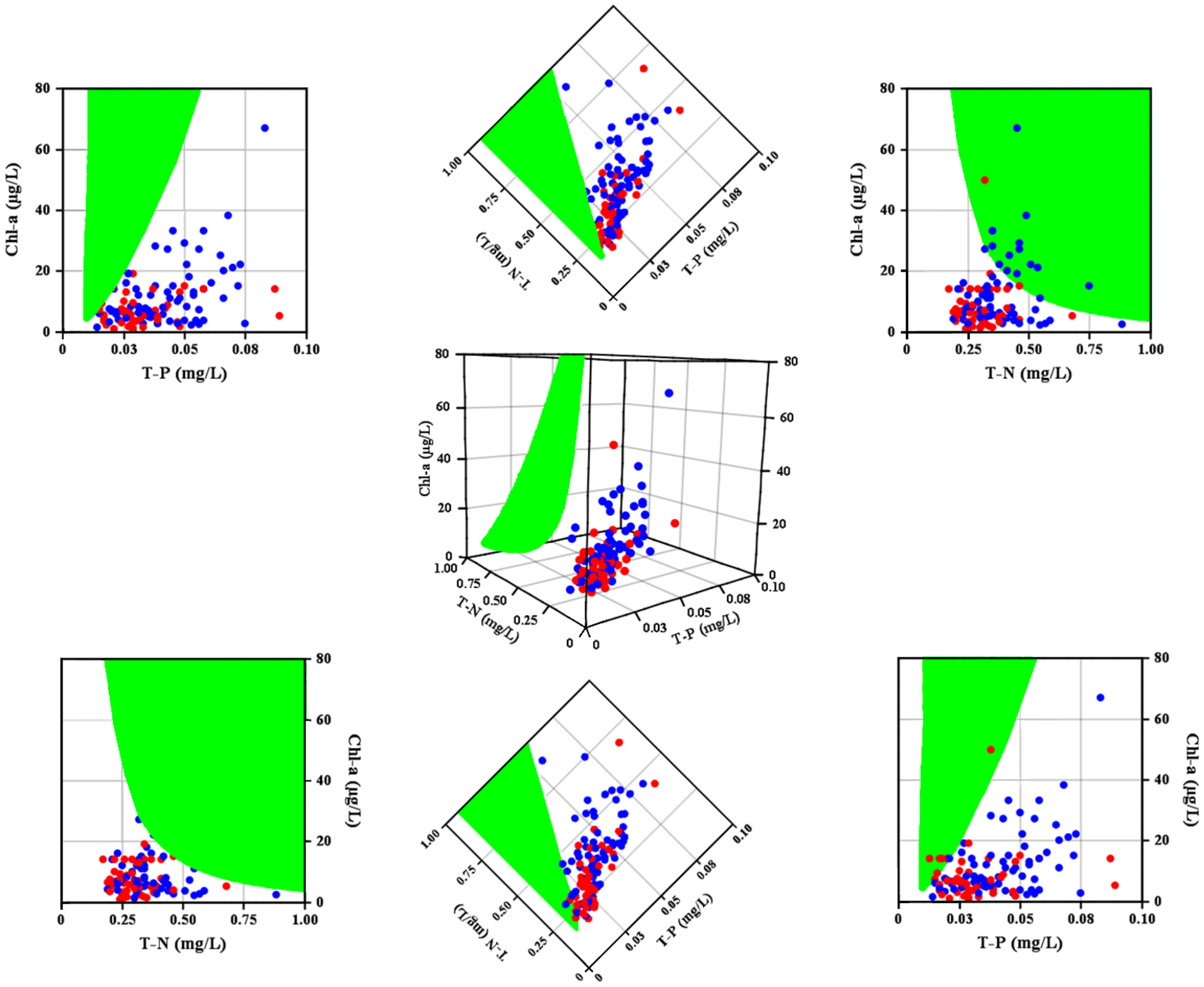


**Figure S4.** Response surface and corresponding 3D scatter plot derived from multiple regression analysis for environmental monitoring site K-7 (fiscal years 2015–2025) (explanatory variables: P and N; response variable: Chl-a). ●: P-addition period; ●: other periods. Projections onto the top and side panels demonstrate that simultaneous increases in P and N in oligotrophic waters—even when following the Redfield ratio—do not necessarily lead to an increase in Chl-a, suggesting the possibility of a different optimal nutrient ratio.

increasing phosphorus (P) and nitrogen (N) concentrations based on the Redfield ratio in oligotrophic coastal waters could paradoxically lead to a decrease in phytoplankton biomass. This suggests that the Redfield ratio is not necessarily optimal for this specific environment. Furthermore, using data compiled from existing literature—with N and P as independent variables and Chl-a as the dependent (response) variable—we estimated the maximum bioavailable concentrations of these nutrients. We then utilized the resulting regression function to perform response surface modeling, constructing the three-dimensional standard response surface shown in **Figure S4**. This standard response surface visualizes changes in Chl-a concentration in relation to N and P levels; in addition to the surface itself, projected views from the top, bottom, and sides are also presented.

**4. Differences in Phosphate Immobilization Mechanisms Depending on Reaction Conditions.**

When excess $FeCl_3 \cdot 6H_2O$ and $KH_2PO_4$ were reacted under acidic conditions (pH < 4, approx. 3.5) and the temporal change in dissolved phosphate concentration was monitored, no precipitate corresponding to solid $FePO_4$ was detected either visually or analytically (**Figure S5**). Under these acidic conditions, excess Fe(III) is expected to exist predominantly as soluble $Fe^{3+}$. However, as observed for Samples 1–4 (**Table 1**), the dissolved phosphate concentration decreased quantitatively in proportion to the dosage of Fe(III). These findings suggest that phosphate immobilization under acidic conditions is driven not by phase transformation into solid $FePO_4$, but rather by the formation of soluble inner-sphere complexes [e.g., $(Fe–H_2PO_4)^+$] or ion pairs between $Fe^{3+}$ and phosphate species (primarily $H_2PO_4^-$). Furthermore, no precipitate formed when $FeCl_3 \cdot 6H_2O$ was reacted with an equimolar amount of $K_2HPO_4$ under acidic conditions. Colorimetric analysis using ammonium molybdate and ascorbic acid revealed almost no free $H_2PO_4^-$, confirming that phosphate forms a robust aqueous complex with Fe(III).

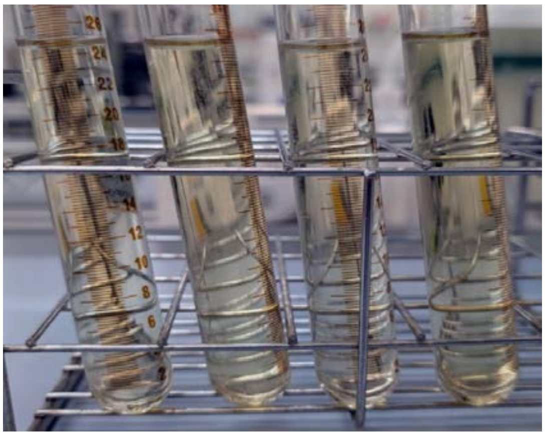

**Figure S5**. Phosphate immobilization by Fe(III) under acidic conditions (pH < 4): **a**. Sample 1, **b**. Sample 2, **c**. Sample 3, **d**. Sample 4.

In contrast, as shown in Samples 5–8 (**Table 1**), under typical seawater conditions (pH approx. 8), Fe(III) predominantly exists as FeOOH, whereas phosphate exists mainly as $HPO_4^{2-}$. Under these conditions, the reaction between Fe(III) and phosphate species is expected to yield $FeOOH–HPO_4^{2-}$ complex precipitates. Indeed, as illustrated in **Figure S6**, while unreacted $HPO_4^{2-}$ and its counter-ion $K^+$

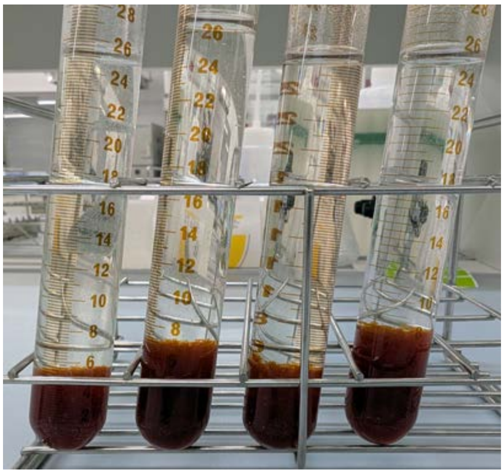

**Figure S6.** Phosphate immobilization by $FeOOH–HPO_4^{2-}$ complexes under neutral conditions (pH ≈ 8): **a**. Sample 5, **b**. Sample 6, **c**. Sample 7, **d**. Sample 8.

remained in the aqueous phase, $Fe^{3+}$ virtually disappeared from the solution due to the precipitation of FeOOH–$HPO_4^{2-}$ complexes and the precipitation of excess $Fe^{3+}$ as FeOOH. A consistent trend was observed when $FeCl_3{\cdot}6H_2O$ was reacted with an equimolar amount of $K_2HPO_4$ at pH approx. 8, resulting in the formation of a precipitate identified as an FeOOH-$HPO_4^{2-}$ complex.

Next, the feasibility of reducing Fe(III) within the generated FeOOH and FeOOH–$HPO_4^{2-}$ complex precipitates to Fe(II) using $NaBH_4$ was evaluated. Upon reduction of the FeOOH precipitate, the resulting $Fe(OH)_2$ was unstable in water and dissolved as $Fe^{2+}$ ions. Conversely, the reduction of Fe(III) within the FeOOH–$HPO_4^{2-}$ complex precipitate led to its dissolution into aqueous $Fe^{2+}$ and $HPO_4^{2-}$ ions. As shown in **Figure S7**, colorimetric analysis using a mixed reagent demonstrated that approximately 100% of the phosphorus in the system was recovered as free phosphate ions ($HPO_4^{2-}$).

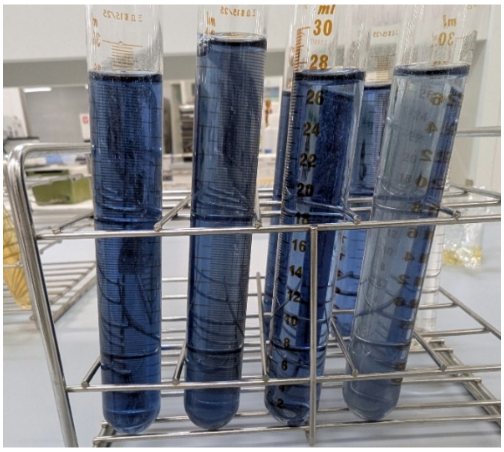

**Figure S7**. Color development using a mixed reagent for free phosphate released into water (reaction under neutral conditions involving the reduction of Fe(III) in FeOOH and FeOOH–$HPO_4^{2-}$ complexes by $NaBH_4$—specifically, color development due to free phosphate derived from the reductive decomposition of FeOOH and FeOOH–$HPO_4^{2-}$ complexes): **a**. Sample 5, **b**. Sample 6, **c**. Sample 7, **d**. Sample 8.
